# Optimization of Experiment Design for Mass Spectrometric Isotopic Labeling Kinetics

**DOI:** 10.1101/331520

**Authors:** Shefali Lathwal, Raaisa Raaisa, Tiago C. Alves, Richard G. Kibbey, Abhishek K. Jha, Graeme F. Mason

## Abstract

Determination of metabolic fluxes by measurement of time-dependent sampling of isotopic enrichments during the administration of labeled substrates provides rich information. Because such experiments are resource-intensive and frequently push the limits of sensitivity of the measurement techniques, optimization of experiment design can improve feasibility with respect to financial and labor costs, time to completion, and increase precision and accuracy of the results. Here we used a previously published set of data acquired in cultured insulinoma cells to evaluate contributions to the sensitivity and variability of the rate of citrate synthase (CS). Specifically, we calculated changes in uncertainty in CS if sample times were dropped or new ones were added, and we observed that some sampling times can be dropped with little effect, while improvements can be made with a strategic choice of when to add samples. We measured the contributions of data sampled at different times on the sensitivity of CS, finding that CS had greater sensitivity at early time points. We tested the concept that if two estimated parameters are correlated significantly, then refining one might constrain the other. In this case, the rate of Beta-oxs was found to be correlated with CS, and narrower variability in Beta-ox did indeed improve the sensitivity of CS. The tests described here might be applied at the initial design stage and then after a pilot phase to improve sensitivities of targeted fluxes and the reduction of materials, time, labor, and other experimental resources. The correlation analyses can be used to consider what orthogonal measurements might be beneficial for further improvement of measurements. While this study used a specific example of a set of time-dependent kinetic isotopic measurements, the results illustrate some generalizable behaviors that can be tested in other experimental systems.

## Introduction

Metabolism has been studied with isotopes for nearly a century, including with stable isotopes [reviewed in (Lehmann, 2017)]. Such kinetic isotopic measurements have the benefit of yielding absolute rates of metabolism, especially when combined with isotopomeric analysis of stable isotopes (Alves et al., 2015; Chance et al., 1983; Gruetter et al., 1994; Katz et al., 1993; Malloy et al., 1987). The derivation of rates and assessment of the limits of understanding depends on mathematical modeling of the metabolic processes under study. Metabolic modeling serves many purposes including testing our understanding of metabolism and quantifying metabolic fluxes. The model is a mathematical expression of our hypotheses about a simplified version of the pathways that we endeavor to study. If the parameters of the model can reasonably be adjusted in a way that the model’s behavior approximates the targeted experimental observables, then our understanding (i.e., hypotheses) may be correct (Garfinkel, 1968). Additionally, if the model approximates the data well, the simulation can yield estimates of the kinetic parameters that govern its behavior, within bounds of uncertainty in those estimates, and it is important that the bounds of uncertainty be known. For metabolic modeling, the uncertainties must often be determined using a Monte Carlo approach (Canavos, 1974; Kuwabara et al., 1990; Mason et al., 1992), as will be explained. Because isotopic metabolic studies are often resource-intensive, in this work we propose a new function of metabolic modeling. Simulations can be used to optimize the experiment design for enhanced sensitivity to particular model features and minimize uncertainties associated with the values of particular metabolic parameters (Mason et al., 1995; Tellinghuisen, 1991).

### Composition of a Metabolic Model

To perform kinetic analyses with time-dependent isotopic labeling studies, several basic components are needed: metabolites (*pools*), substrate sources (*drivers*), metabolic reactions that carry isotope from one metabolite to another (*rates*), and the measured isotopic labeling data (*targets)*. The pools, drivers, and rates are combined to create a set of mass and isotope balance equations, with one of each for every relevant labeling pattern of each metabolic pool that is in the model.

The mass balance equations are mathematical statements of how the total concentration of a metabolic pool (‘mass’ referring to its size, not to be confused with its molecular weight) changes over time. A mass balance equation has the general form

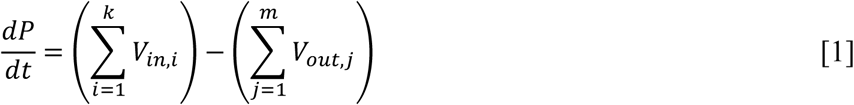

where 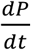 is the rate of change of the mass of the metabolic pool, *V*_*in,i*_ is the rate of the *i*^th^ flow into the pool of metabolite *P*, and *V*_*out,j*_ is the rate of the *j*^th^ flow out of the pool. Many isotopic labeling measurements are performed at a metabolic steady state, so 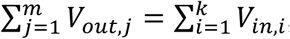 and consequently the size of the pool *P* does not change with time. However, in kinetic isotope studies the isotopic labeling changes with time, and the isotope balance equations show quantitatively how the isotopically labeled fraction of a metabolic pool changes, with the general form:

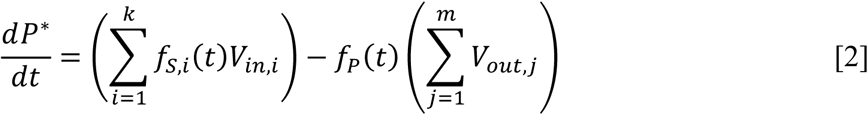

where *S, i* is the i_th_ of k substrates that provide mass for the pool, each at a rate *V*_*in,i*_ *f* _*S,i*_ is the fraction of the i_th_ substrate that is labeled at any instant *t, f*_*P*_ is the fraction of the product *P\** that is labeled at the instant *t*, and is the rate of the ^th^ flow out of the pool. Because ^13^C has been shown to have a negligible isotope effect (Attwood et al., 1986; Melzer and Schmidt, 1987; Tipton and Cleland, 1988), the rate of isotopic inflow from each source is equal to the rate of total mass flow from that source, multiplied by the fraction of the flow that is ^13^C-labeled. The mass and isotope balance equations can then be solved in light of various constraints that are placed on the model by mass balance and a priori knowledge from one’s own experiments and reliable literature and other sources.

Once a metabolic model is created that is able to achieve a reasonable approximation to the data, one can begin to approach issues of experimental design in a quantitative way. Isotopic labeling studies are usually intensive with respect to time, labor, and materials, and it is common for some parts of measurements to approach the limits of sensitivity of the methods of detection. Quantitative experimental design considerations have the potential to reduce costs, improve sensitivity of the values of metabolic parameters to the data, and increase throughput given limited time, money, and resources. Although estimates of rates are the most common objectives for kinetic fitting, such data can and has sometimes been used to measure other parameters, such as the concentrations of metabolites in specific cellular compartments and enzyme kinetic constants (Patel et al., 2010).

### Assessment of Parametric Uncertainties

A key aspect of quantitative understanding of a system is the knowledge of how precisely the system’s parameters are known and how much the imprecision can impact the calculated outcome. To this end, an analysis of uncertainty is needed (Kennedy and O’Hagan, 2001). In many experiments, group statistics can be used to ask, for example, what is the standard deviation of the height of students who receive vitamin supplements? However, unlike in a group of students, where each individual yields one value for height, in isotopic kinetic studies, many individual measurements are needed to be analyzed as a group to yield one value of each metabolic parameter, so there is no list of rates for individuals to provide group statistics like a mean and standard deviation. Furthermore, the scatter in the data can be an appreciable fraction of the signal. With relatively large scatter and approximate exponential behavior, the uncertainties (i.e., the probability distributions) of the estimated parameters often do not follow Gaussian distributions. Hence, a Monte Carlo approach can be used to assess the effects of noise in these scenarios.

If we consider a metabolite with constant inflow of label and steady state conditions, it can be modeled by the differential equation:

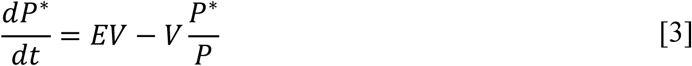

Here, *P\** is the labeled pool of metabolite, *P* the total pool, *V=V*_*in*_ *= V* _out_ is the rate of inflow of label. The solution has the form:

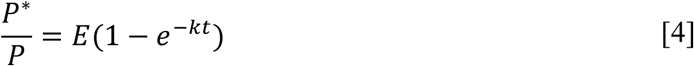

where *E* is the enrichment of the source and.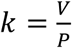 We use this model to explore the impact of noise on distribution of estimated values of *k* with *E* = 1 and *k* = 0.1 (Mason and Rothman, 2004). Measurements of *P\** would, with noise, resemble Fig. 1A, and the scatter of the measurements would force the estimated exponential rate constant k to have a range of uncertainty. Monte Carlo analysis is performed by first treating the least-squares, fitted curve as the best estimate of the true behavior of the system, and then adding randomly distributed, representative noise repeatedly to that best estimate. New fits are then calculated to each of the simulations that can be plotted as histograms. This procedure effectively asks what the distribution of values would be if the same experiment were performed repeatedly under the same conditions. For the example in Fig. 1, 15,000 simulations were run with noise levels of 5%, 10%, and 20%, and the resulting distributions of the rate constant k (Fig. 1B) show that as the level of the noise increases, the distribution broadens and becomes increasingly non-normal.

**Figure 1:**
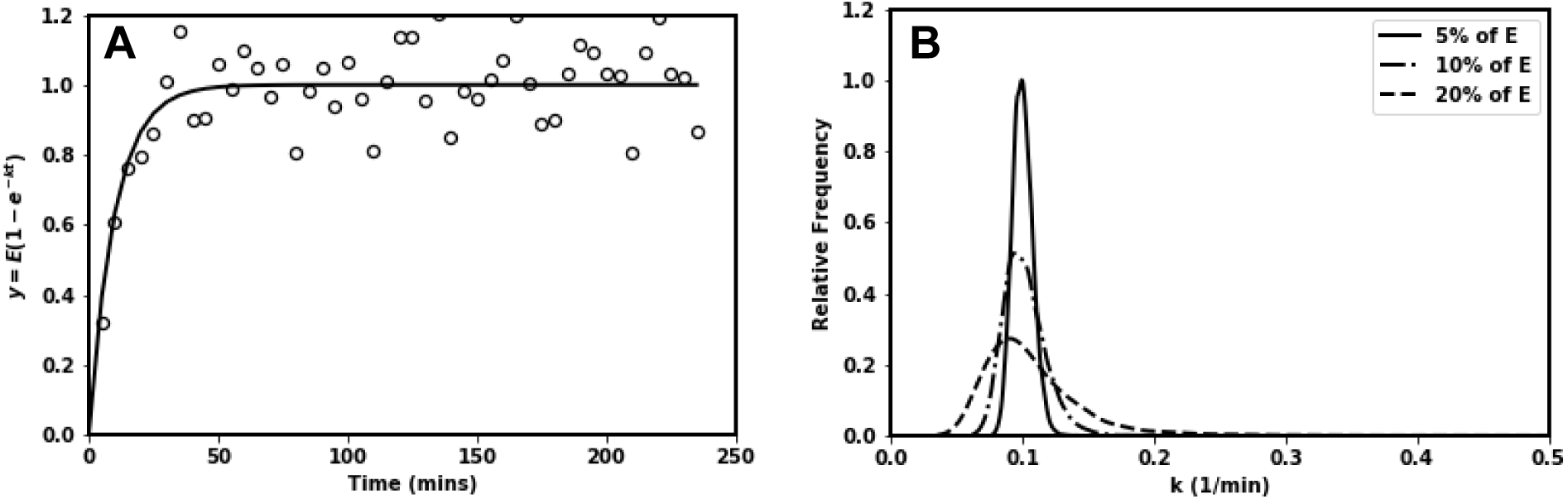
Monte Carlo analysis of fitting to an exponential curve, y = E(1 - e^-kt^), by iteration of the values of E and k. (A) Representation of data with 10% noise corresponding to an exponential curve with E=1 and k=0.1. (B) Distribution of fitted rate constant ‘k’ for three levels of noise, 5%, 10% and 20% of the value of E.

### Overview of Procedures

A published data set from insulinoma cells treated with [U-_13_C_6_]-D-glucose at four different concentrations (2.5-9 mM) and seven different time points (0-240 min) and central carbon metabolites analyzed by LC-MS/MS were used as source data for the simulations (Alves et al., 2015). In the original manuscript, several metabolic rates were calculated from the data using the kinetic model of Fig. 2. Here, experiment sensitivity and efficiency were explored using those data and the kinetic model. In particular, citrate synthase (CS) was examined because it is central to understanding all of the elements of the TCA cycle and consequently a key element with respect to cellular energetics, mitochondrial health, and other biomedically related issues. A secondary focus were the rates of beta oxidation of fatty acids (Beta-ox) because of their impact on oxidative substrate selection. The sensitivity of the values of CS and Beta-ox to the timing of sampling was investigated, considering whether experimental precision might be enhanced by judicious sampling and whether some time samples could be eliminated with little effect on precision.

**Figure 2:**
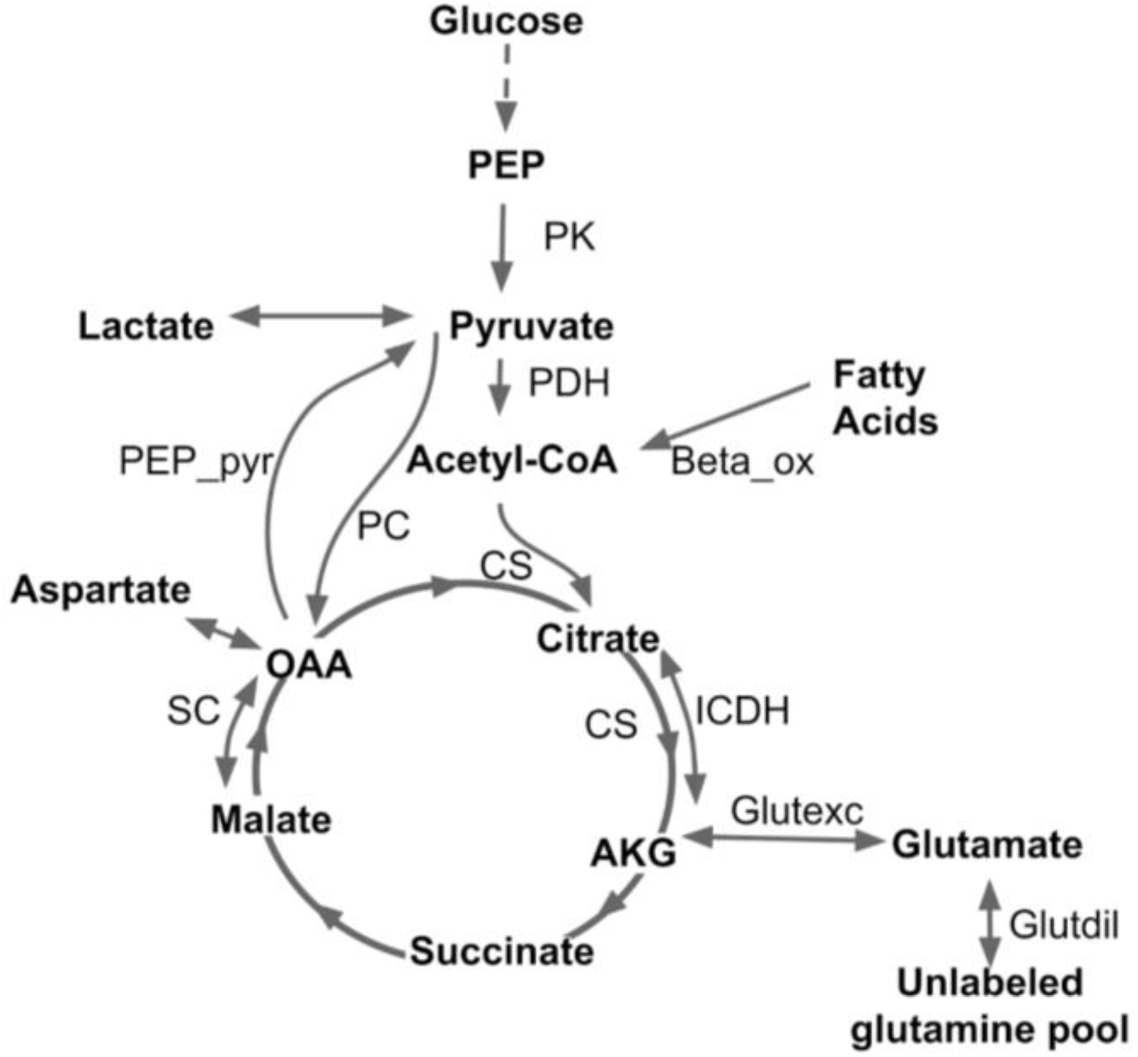
Representation of the TCA cycle metabolites and fluxes used in the model (Alves et al., 2015). Rates and Concentrations are given in Table 1 and Table 2 respectively.

## Methods

### Kinetic modeling for flux analysis

The source data (Alves et al., 2015) consisted of central carbon metabolites (phosphoenolpyruvate (PEP), pyruvate, lactate, citrate, glutamate, succinate, malate, and aspartate) from an insulinoma cell line INS-1 832/13 at a range of time following incubation with [U-_13_C_6_]-D-glucose. Positional enrichments were calculated as described using Mass Isotopomer Multi-Ordinate Spectral Analysis (MIMOSA). The enrichments of pyruvate, citrate, glutamate, succinate, and malate were used as target data to be fitted with the differential equations that describe the pathway shown in Fig. 2 (Alves et al., 2015). The fitting was performed using a python kinetic flux package, PollyPhi **™**Absolute (Elucidata, Inc.) described in Supplemental Material S1. Two optimization algorithms, sequential least squares programming (SLSQP) and least squares, were used to fit the data and obtain the values of several fluxes including: citrate synthase (CS), beta oxidation of fatty acids (Beta-ox), exchange between citrate/isocitrate and alpha-ketoglutarate (αKG) (ICDH), exchange between malate/fumarate and OAA (SC), exchange between αKG and Glutamate (Glutexc), dilution of glutamate (Glutdil), and pyruvate carboxylase (PC). The flux values were expressed in terms of μM/μM Taurine/min.

**Table 2:** Concentrations of metabolites used in the model

| Glucose<br>(mM) | Concentration ($\mu\text{M}/\mu\text{M}$ Taurine) | | | | | | | | |
| --- | --- | --- | --- | --- | --- | --- | --- | --- | --- |
| | Succinate | Pyruvate | Citrate | Glutamate | Malate | PEP | OAA | AcetylCoA | $\alpha\text{KG}$ |
| 2.5 | 0.020 | 0.004 | 0.219 | 3.949 | 0.380 | 0.024 | 0.020 | 0.020 | 0.020 |
| 5 | 0.015 | 0.014 | 0.27 | 4.26 | 0.37 | 0.023 | 0.015 | 0.015 | 0.015 |
| 7 | 0.0180 | 0.0353 | 0.47 | 5.09 | 0.94 | 0.0564 | 0.0180 | 0.0180 | 0.0180 |
| 9 | 0.0399 | 0.0697 | 1.31 | 7.63 | 2.02 | 0.1137 | 0.0399 | 0.0399 | 0.0399 |

**Table 1:** Absolute flux values obtained from fits in PollyPhi ™ Absolute

| Glucose<br>(mM) | Fluxes ($\mu\text{M}/\mu\text{M}$ Taurine/min) | | | | | | |
| --- | --- | --- | --- | --- | --- | --- | --- |
|  | CS | Beta-ox | PC | ICDH | Glutdil | Glutexc | SC |
| 2.5 | 0.106±0.008 | 0.016±0.000 | 0.002±0.000 | 0.062±0.009 | 0.302±0.030 | 7.43±0.5 | 7.21±0.5 |
| 5 | 0.089±0.002 | 0.011±0.000 | 0.005±0.000 | 0.041±0.004 | 0.105±0.003 | 20.2±2.2 | 20.3±2.0 |
| 7 | 0.131±0.003 | 0.021±0.003 | 0.021±0.001 | 0.030±0.005 | 0.139±0.005 | 36.8±4.5 | 37.8±4.8 |
| 9 | 0.234±0.004 | 0 | 0.050±0.001 | 0.124±0.005 | 0.268±0.010 | 62.8±7.1 | 42.0±1.3 |

### Assessment of timing of data acquisition on uncertainty distributions of estimated parameters

Each target species used in the objective function had data at seven time points, with six replicate measurements at each time point. To perform Monte Carlo analyses, the replicates were assumed to be part of a normal distribution with a mean and standard deviation equal to the mean and standard deviation, respectively, of the measured data for each target at each time point. These normal distributions were then randomly sampled 500 times to generate new, artificially noisy datasets. Any negative values in these datasets were replaced with zero, just as is done with actual enrichment data. Each of these new datasets were fitted with the model independently to create a list of 500 different parametric fits and estimate the distribution and correlative relationships of flux values.

### Simulating an experiment with data collected at fewer time points

Data sets with fewer time points were simulated by removing individual time points from the experimental target. Monte Carlo simulations were also run with each modified data set to obtain new distributions of fluxes. The purpose was to assess if a more efficient experiment could have been performed, one that yielded acceptable precision for the value of CS but with fewer data points.

### Simulating an experiment with data collected at more time points

To assess additional data sampling might be improve the parameter estimations, simulated data were generated along the original least-squares fit by creating 15 time points spaced uniformly between 0 and 50 minutes. Here, the goal was to assess how data at any given time point impacted the value of CS. These simulated points were combined with the experimental data measured at 60, 120, and 240 minutes, effectively simulating how the precision of the value of CS might improve if additional data were acquired at early time points. The decision to focus on the early time points arose because at early time points, the enrichments are changing more quickly and the kinetics have more impact than later, as the metabolites’ labeling approach a steady state.

### Comparison of distributions achieved with different experiment designs

The distribution of parameters obtained after Monte Carlo analysis was either plotted directly as a histogram or fitted to a beta distribution for visualization. To quantify the uncertainty in a given parameter, the distribution of values of that parameter was randomly sampled to make 50 groups of 20 values each. The standard deviation of each group was then calculated and the average of these 50 standard deviations was used as a quantitative measure of parameter uncertainty.

### Determination of sensitivity of parameters to specific components of the data

A sensitivity analysis was performed to determine which targets and specific time points have the largest effects on the estimated flux parameters. After the best-fit estimates of the fluxes were obtained from the experimental data, the sensitivities of those fitted parameters to the data were estimated by the following procedure: the time courses of each labeled species obtained with the estimated fluxes were assumed to represent the true behavior of the system. Then, a fixed positive deviation was introduced in the calculated time course of a given target at a given time point, and the optimization algorithm was run again. This procedure effectively poses the question, “If the data were higher at a particular time point, how much would it change the estimated fluxes?” These steps were repeated for each target at each of the seven time points one-by-one to obtain new flux values. Sensitivity of a flux was then calculated as

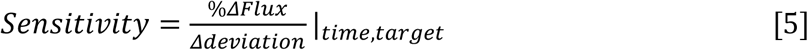

### Correlation analysis from Monte Carlo lists

The 7 mM glucose group was chosen to illustrate the correlations between fluxes because it showed a broader distribution of uncertainty for Beta-ox than did the 9 mM group. Five hundred Monte Carlo simulations were performed with and without the 240-minute time point. The resulting distributions from these simulations were used to calculate Pearson correlation coefficients for pairwise comparison of fluxes.

## Results

The least-squares fits for the 7mM data set are shown in Fig. 3A, and the resulting flux values are given in Table 1. After least-squares fitting, the distributions of uncertainty for CS and Beta-ox were estimated with Monte Carlo iterations consisting of 500 runs per data set. These distributions showed a non-normal distribution (Fig. 3B, 3C), as expected from noisy datasets (Fig. 1B).

**Figure 3:**
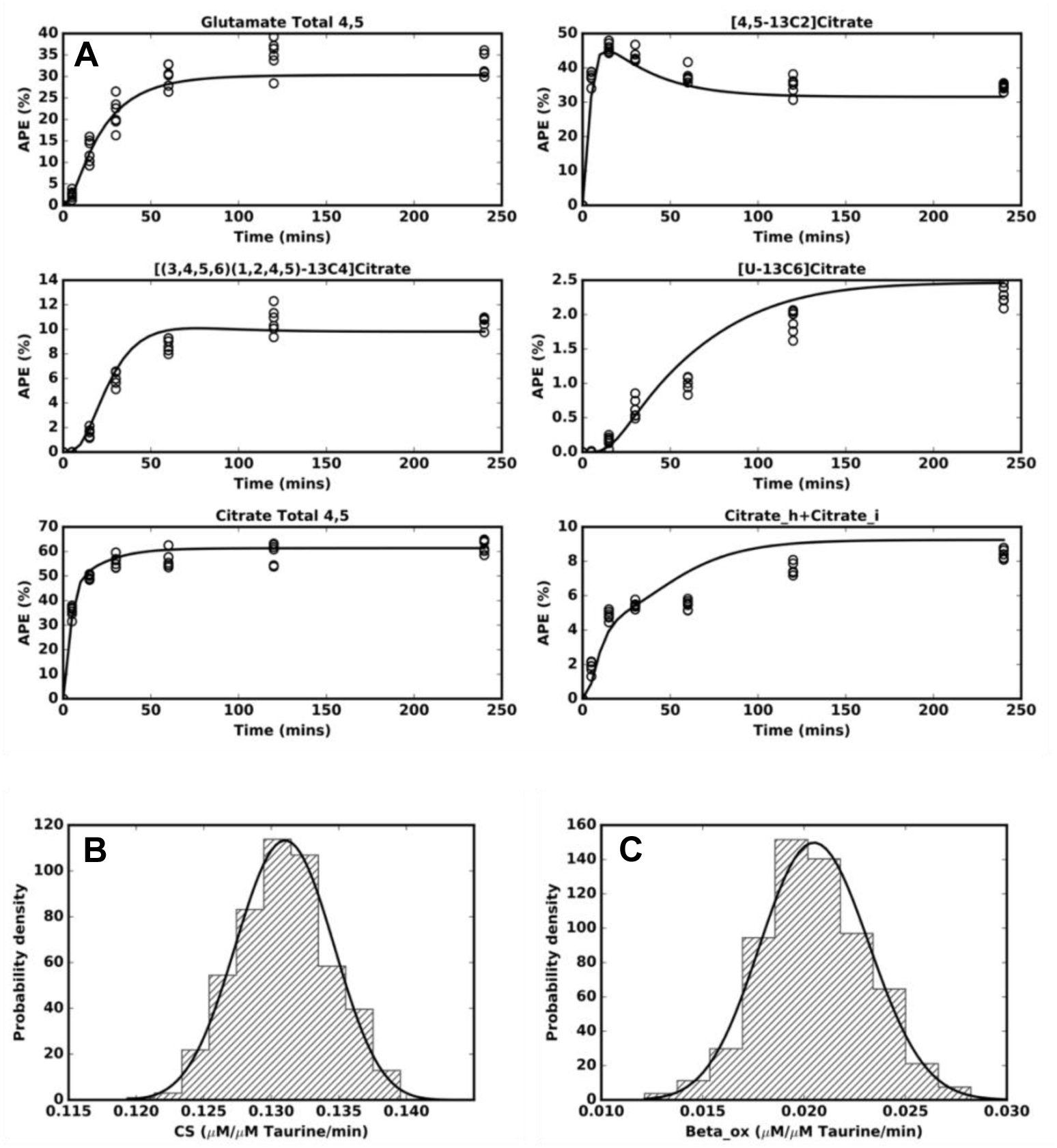
Fit and distribution of fitted parameters. (A) Least-square fits (solid lines) of the model of Figure 2 to the experimental data (open circles) for the data acquired with 7mM glucose. Glutamate Total 4,5 is the sum of all isotopomers of glutamate which have are labeled in carbon positions 4 and 5, Citrate Total 4,5 is the sum of all isotopomers of citrate which have a label in positions 4 and 5, and Citrate_h+Citrate_i refers to the combination of [(1,2,3,4,5)(2,3,4,5,6)(1,2,4,5,6) and (1,3,4,5,6)-_13_C_5_]Citrate. Distributions of (B) CS, and (C) Beta-ox obtained from 500 Monte Carlo simulations. The histograms have been normalized such that the shaded area is equal to one.

### Time-dependent sensitivity with MIMOSA sampling

The heatmap in Fig. 4 shows the sensitivities of the estimated CS to the target data for each measured species (vertical axis) over the range of time points (horizontal axis), for the 9 mM glucose condition. The values in the heatmap represent the percentage change in the estimated value of CS if the enrichment of a given target at a given time point were higher than that measured in the experiment. The magnitude of the sensitivity varies with both the target species and time.

**Figure 4:**
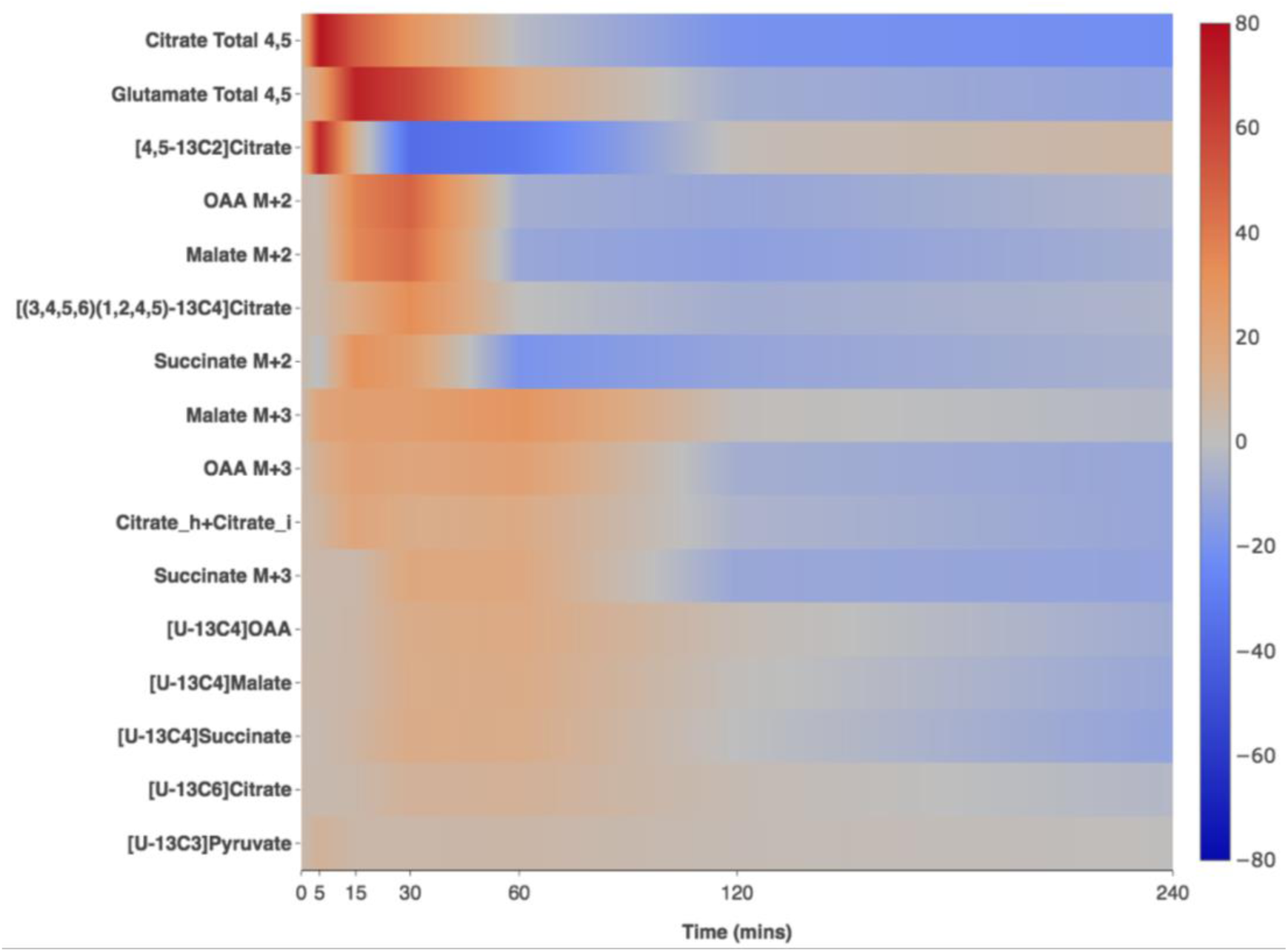
Percentage sensitivity of the estimated CS flux (9mM glucose condition) to the experimental targets and time. The colors represent percentage change in CS caused by deviations in the enrichment data, with greater color saturation representing large responses of CS.

With respect to target species, the highest sensitivities to the CS estimate were 77%, 72%, and 71%, observed for Citrate Total 4,5, Glutamate Total 4,5, and [4,5-_13_C_2_] Citrate, respectively, indicating that tighter measurements of Citrate and Glutamate can lead to tighter CS estimates. For many species, especially the ones that have low steady state enrichment, sensitivity of CS was close to zero at all time points, which means that in general metabolites with low labeling have little impact on the value of CS. Therefore, the estimation of CS is generally insensitive to target species that are less labeled, even though they have lower signal-to-noise ratios.

For time-dependence, the highest sensitivities observed for any target at 5, 15, 30, 60, 120, and 240 minutes were 77%, 72%, 59%, 29%, 3%, 7%, respectively. These values indicate that individual targets at early time points (up to 30 minutes) can have a large impact on the estimated values of CS and at later time points, have less effect. The time-dependent sensitivities also illustrate that for kinetic studies, the majority of the information is captured when the enrichments are changing most rapidly. Since the sensitivity values are calculated for each target species used in the objective function, they will change if the target data are changed.

### Effect of altered experimental sampling

We eliminated data at all time points one by one to quantify the effect of each time on the distribution of the estimated value of CS (Fig. 5, S1, S2, and S3). The data at a given time could have an effect on both the mean value and the spread of the distribution (i.e., the tightness of the estimated CS). We observed that for 9mM glucose concentration, omission of data at any time point, except 60 and 120 minutes, led to a statistically significant change in the distribution of CS. In other words, if only six time points were measured at either 0, 5, 15, 30, 120, and 240 minutes or 0, 5, 15, 30, 60, and 240 minutes, the distribution of CS would have been statistically similar to when all seven time points, 0, 5, 15, 30, 60, 120 and 240 minutes, were measured. As seen from the sensitivity map in Fig. 4, early time points, 5 and 15 minutes led to maximum change in the expected values of CS (Fig. 5A). We also observed that once an isotopic steady state is approached, only one time after that needs to be sampled, but it is crucial to have that late sample. For example, without the data at 120 minutes, but in the presence of data at 240 minutes, the distribution of CS was statistically the same as when the data at both 120 and 240 minutes was present. However, in the absence of data at 240 minutes, but presence of data at 120 minutes (where some enrichments have not reached an isotopic steady state), the distributions of CS differ (Fig. 5B). If data at both 120 and 240 minutes are absent, the distribution of CS changed significantly compared to the case when all time points have been included. The effect of removing the 240-minute time point on CS is prominent in the estimated CS for 5mM glucose concentration as well (Fig. S2). As seen in Fig. 5, the removal of a single time point in some cases can increase the uncertainty of CS (15 minutes). Another factor to consider is the potential for bias in the expected value, which was seen at some of the time points. Removal of data at 5, 15, 30, or 240 minutes led to a statistically significant shift in the expected value of CS, which reflects the potential for bias in the values of fitted parameters when insufficient data are acquired. Taken together, these two points suggest that if the removal of a time point changes neither the uncertainty nor the expected value of the parameters of interest, the point is unnecessary. In other cases, in which either the distribution or the expected value changes, the point is needed.

**Figure 5.**
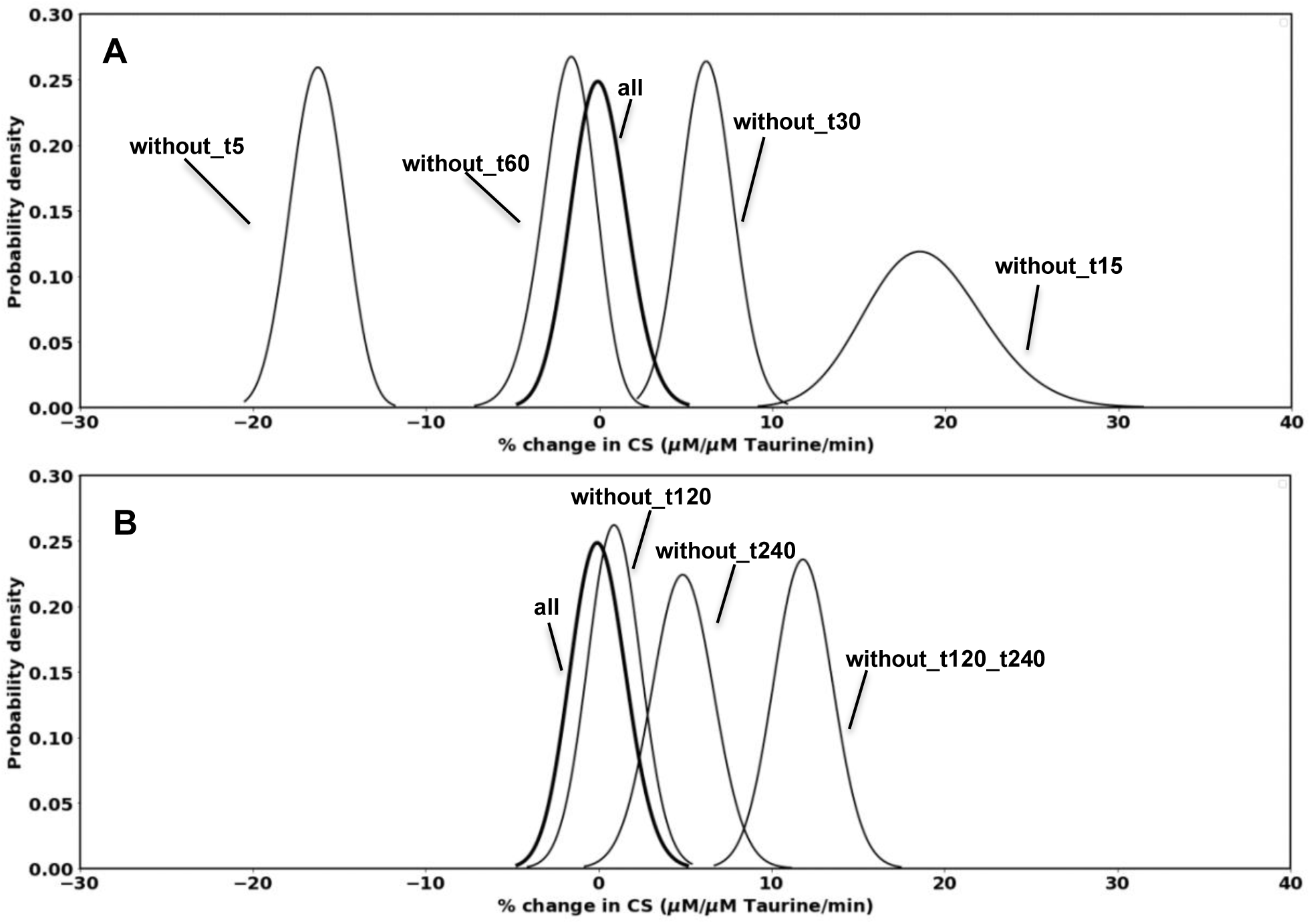
Distributions of CS (9mM glucose condition) relative to the value obtained when all time points are used or some are eliminated. The x-axis represents percentage change in CS compared to the average value for the condition where all seven time points (0, 5, 15, 30, 60, 120 and 240 minutes) have been used. Note that the curve “all” is centered around zero because its expected value is the reference for the test cases. The Supplemental Material shows the corresponding plots for data acquired with 2.5, 5, and 7 mM.

While the analysis of the importance of each time point is useful in retrospect, we recognize that the experiment design stage requires general rules of thumb to determine a range of time points that are likely to be important for sampling. To do this, we first defined a representative rate constant *k* for each of the four datasets in our study as the ratio of the primary flux through the pathway divided by the concentration of the largest metabolite pool (i.e., *flux*/*concentration*), and the rate constants were used to estimate labeling half-times *t*_*1/2*_ (Supplementary Table S1). If we approximate the time courses with these rate constants and compare the time of sampling (Fig. S4) used by Alves et al., we find that the experimental measurements are well distributed over the entire dynamic range for all four datasets. We expand on this concept later in the Discussion, recommending a procedure to choose sample times when planning new flux experiments.

### Tightening precision of selected parameters

When there is a strong correlation between fluxes in a pathway, it suggests that if we reduce the uncertainty in any one of these fluxes, the uncertainty in the other correlated fluxes should also decrease. We used Beta-ox and CS as examples because these fluxes are strongly correlated as evidenced from a comparison of values obtained from the 500 Monte Carlo simulations for the 7mM glucose dataset (Fig. 6). We tested our hypothesis by comparing the distribution of CS and Beta-ox between two conditions: 1) data collected until the steady state is reached for all isotopomers (i.e., last data point collected at 240 minutes), and 2) data collection stopped before the steady state is fully established (i.e., last data point collected at 120 minutes). We observed that the uncertainty in both CS and Beta-ox was lower in condition 1 compared to condition 2. The lower uncertainty was because a clearly defined steady state limits the possible values of fluxes and so more strongly constrains the optimization algorithm. It is worth noting that the inclusion of the 240 minute point tightened the uncertainty in Beta-ox more than the uncertainty in CS, reflecting the fact that the 240 minute point contains more information about Beta-ox than it does about CS.

**Figure 6:**
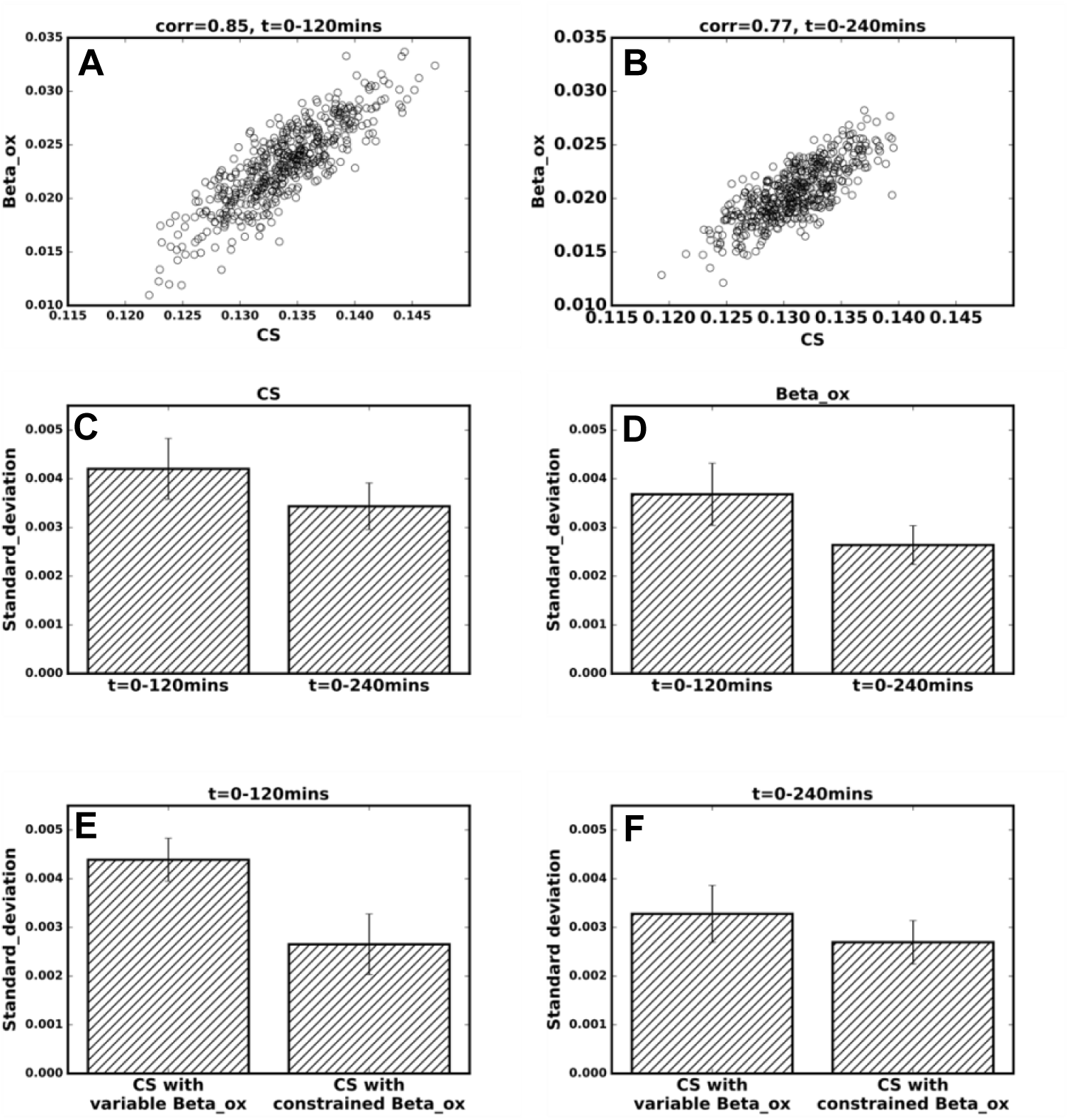
Correlation between CS and Beta-ox and the effect of their correlation on their uncertainty under different conditions. Correlation between estimated values of CS and Beta-ox obtained from Monte Carlo simulation of 7mM glucose condition with data considered from (A) 0-120 minutes, and (B) 0-240 minutes. Comparison between standard deviation in the estimated values of (C) CS and (D) Beta-ox considering data from 0-120 minutes and 0-240 minutes. Comparison between the uncertainty in CS with unconstrained and constrained Beta-ox for the conditions where data is considered from (E) 0-120 minutes, and (F) 0-240 minutes.

As shown in Fig. 6E and F, we also compared the distributions of Beta-ox and CS under the following conditions: 1) both Beta-ox and CS were allowed to vary and settle to a best-fit estimate, and 2) the value of Beta-ox was constrained to a fraction of CS based on the relative mass isotopomer distributions of pyruvate and acetyl CoA at 240 minutes. Compared to the case with both Beta-ox and CS varied freely, when Beta-ox was constrained, the uncertainty in CS was reduced. This exercise shows that if the relative value of Beta-ox can be determined with great precision, then the value of CS or other parameters correlated with the value of Beta-ox will be known more precisely. This test was repeated for experimental data limited to times up to 120 minutes, and constraining Beta-ox led to an even greater reduction in the uncertainty of CS. The 240 minute data contain a great deal of information about Beta-ox relative to CS, so the additional constraint achieved by fixing the relative rate of Beta-ox contributes only a small amount to the precision of CS (Fig. 6F). When the 240 minute point is not used, then Beta-ox is less well determined, so the benefit of constraining the value of Beta-ox is greater (Fig. 6E). Practically, this means that one could eliminate the 240 minute point if there were an independent way to assess Beta-ox.

## Discussion

A common approach for quantifying metabolic fluxes in a pathway involves the use of stable isotopic tracers, with subsequent labeling patterns measured at multiple time points. Once the data have been collected, biological insights can only be derived after the data have been processed for metabolites or their fragments, corrected for natural abundance of the isotopic tracer and a model has been fitted to the data to estimate the values of fluxes. Since we do not have a direct experimental measure of the fluxes, it is important to determine the statistical significance of the estimated fluxes and to identify the experimental variables which have the largest effect on the estimated fluxes and the extent of those effects.

Some common experimental variables that can affect the accuracy of the estimated fluxes are, 1) the number of time points at which data are collected, 2) the placement of the time points being measured, 3) the number of replicates at a given time point, 4) any variables or additional measurements available to constrain the model, and 5) the information present in the collected data (e.g., isotopologue information that is present in LC-MS data or positional (isotopomer) information of the labeling patterns that is present in LC-MS/MS data and/or Nuclear Magnetic Resonance (NMR) data). It has recently been shown that the information about the position of the label in LC-MS/MS data led to more accurate flux estimates compared to only mass isotopologue information present in the LC-MS data (Alves et al., 2015). The spread of the distribution in flux through PC decreased by about half when isotopomer data were used instead of isotopologue data. In the present work, we quantified the effect of experimentally measured targets, time points at which data have been collected, and the correlation between parameters on the estimated fluxes of the TCA cycle in insulinoma cells (Alves et al., 2015).

Fig. 7 shows a suggested procedure for piloting and refining experimental sampling, including number of samples and their timing to cover a range of potential metabolic rates to meet required limits of precision. As we examined the effects of parametric sensitivity to time, we observed that certain parameters like flux through CS were defined more precisely by inclusion of some time points, such as clear establishment of an isotopic steady state, and capturing the rapid change in enrichments at the beginning of the experiment. Therefore, the time points to be measured for any given pathway should be distributed such that most, but not all, of the data are collected at the beginning, where the label distributions are changing rapidly. It is also important to collect some data to establish that an isotopic steady state has been reached. Since the time scale of any metabolic process depends on many factors (e.g., the labeled nutrient being used, the type of cell, the state of the cell, and the pathway being studied), the optimized selection of time points requires an estimate of the time scale of the pathway under investigation. As discussed in Results, the labeling times can be estimated from a range of fluxes and metabolite concentrations expected from published values or pilot measurements. If one obtains upper and lower bounds of the labeling half-times *t*_*1/2min*_ and *t*_*1/2max*_, time points can be selected strategically as prescribed in Fig. 7 to achieve accurate kinetic measurements over an anticipated range of rates. For piloting, it is often beneficial to begin experimenting with the early estimated time points and proceed sequentially through measurements to the later sample times. The benefit of sequential measurements is that the experimenter may discover that the isotopic steady state has been established earlier than would be the case for the slowest possible time constant, rendering the longest suggested time points unnecessary. One must also consider physiological constraints, such as avoiding too long an experiment because of possible undesirable interference due to other cellular processes that occur at longer time scales.

**Figure 7:**
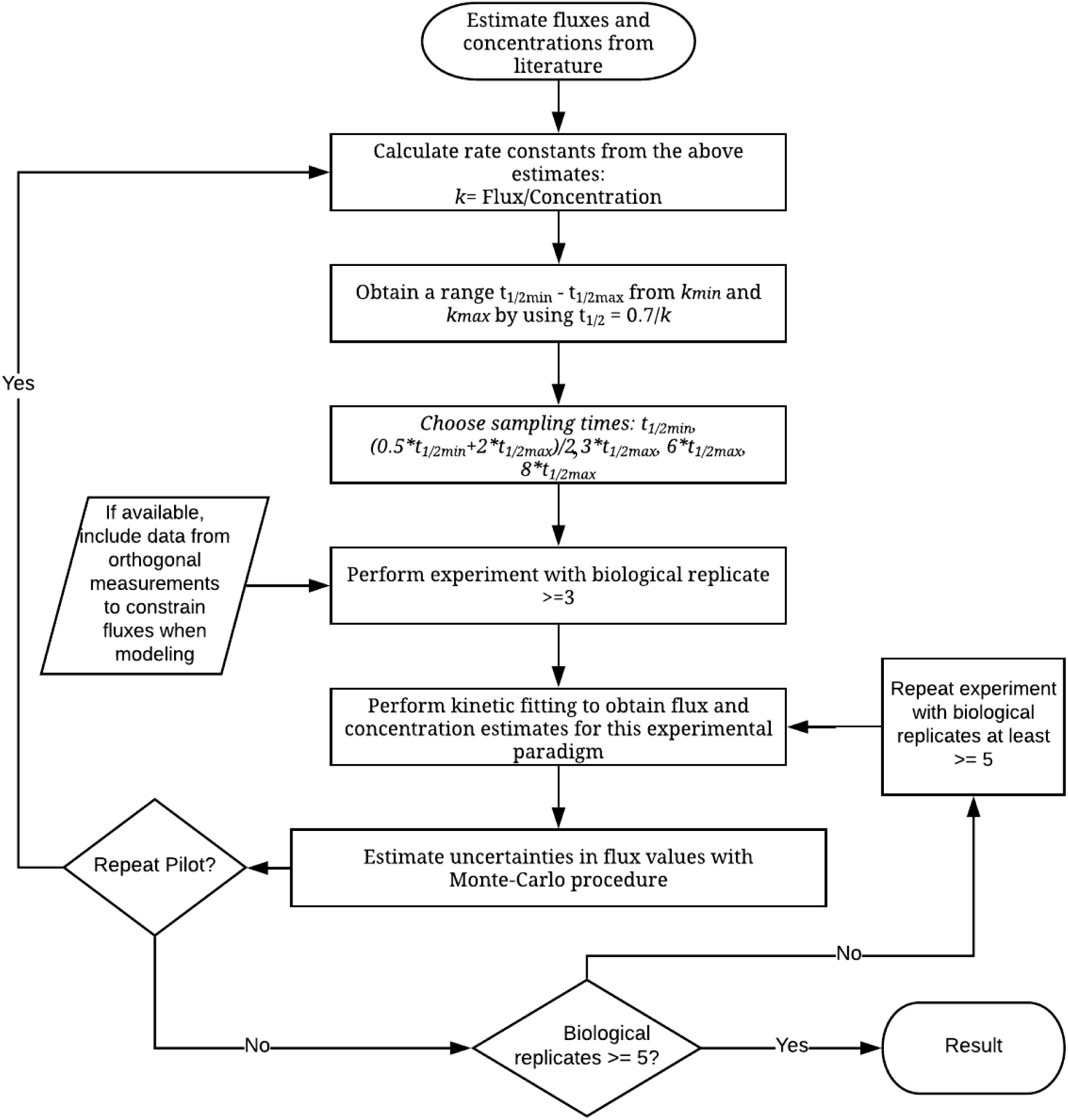
A flowchart of suggested steps for designing and refining an experiment. Initial estimates of fluxes and concentrations, often from literature, can be used to approximate upper and lower labeling half-times *t*_*1/2min*_ and *t*_*1/2max*_, using the formula *t*_*1/2*_ = ln(2)/*k* ∼ 0.7/*k*. The *t*_*1/2min*_ and *t*_*1/2max*_ are used to select specific sampling times as demonstrated in Fig. S5. Pilot experiments with three or more biological replicates are recommended to support statistical approximation for estimation of actual *t*_*1/2*_ values and refinement of the time points if needed. If feasible, data from orthogonal measurements (e.g., Beta-ox) may constrain fluxes in the kinetic analysis. Generally, it is necessary to perform additional measurements to achieve at least 5-6 replicates at each time point, to perform a more based on required precision for the biological questions under consideration.

We found that the values of some parameters were correlated, and so we hypothesized that improving the precision of a few parameters would allow us to reduce the uncertainty in the values of other correlated parameters. We tested this idea for the values of CS and Beta-ox and showed that the correlation between these fluxes meant that constraining the value of Beta-ox led to a tighter distribution of the estimated CS. This approach is routinely used for metabolic studies of brain, where relative rates are constrained using steady-state measurements obtained with one isotope while dynamic rates are assessed with another (Patel et al., 2005). We propose that in addition to using the labeling data at steady state, any other measurements (such as nutrient uptake or secretion rates) that can be used to constrain fluxes, should when possible be included in the experiment design. Another type of orthogonal restraint is the calculation of some of the same fluxes using different labeled substrates.

While we focused the present study around CS and Beta-ox, it is important to note that other fluxes in the pathway could behave in somewhat different ways but would be expected to follow similar principles. The size of the noise or scatter of the data also affects the shape and width of the distributions of uncertainty (Fig. 1B), but that was not directly investigated in the present work. For noisy data, uncertainty distributions can be non-normal and the median may be better than the mean to approximate the best-fit value (Hooker, 1907). It is recommended that biological replicates should be used at each measured time point to get an estimate of noise in the targets and determine its effect. It may become clear that the noise must be reduced if the experiment is to yield the required precision.

We also observed that if the time points are not distributed optimally, then not only the spread in the estimated values of the flux (uncertainty) will be affected, but that also the estimates of fluxes can be biased up or down. Such bias will not be captured in routine statistical analysis of flux estimates, but it should be considered while designing the experiment. Biases can also be introduced if there are errors in the recording of sampling times. For example, if a time point is recorded as six minutes after the start of isotope exposure but was actually captured at five minutes, the estimated fluxes can be significantly overestimated (Supplementary Material S6).

The insights derived in this work are summarized in Fig. 7 and can be applied to reduce the use of experimental resources by eliminating acquisition of data that contribute minimally to accuracy and precision of the estimated fluxes. The results demonstrate that the fitting incorporates results from independent measurements, such as one might obtain with different isotopic labeling studies or with other, non-isotopic techniques, can constrain the fitting and further enhance the accuracy and precision of the estimated fluxes. Judicious sampling and strategic addition of information can improve the feasibility of time-course labeling experiments, which are often time and cost intensive due to the price of reagents, large number of samples, and the expertise required in sample preparation and data collection. The benefits will vary according to the limitations of each experiment. Some experiments may be limited by budget constraints, whereas others may be driven primarily by a need for quick results or high precision. We hope that by highlighting the effect of experimental variables on the estimated flux values, we have given a set of tools to help researchers in judicious experiment design where they can balance labor, use of materials, costs, and time according to their specific experimental needs.

**Author Contributions**
Shefali Lathwal designed PollyPhi Absolute(R), ran calculations, wrote manuscript. Raaisa designed PollyPhi Absolute(R), ran calculations, wrote manuscript. Tiago C. Alves acquired mass spectrometric data, contributed to manuscript. Richard Kibbey oversaw acquisition of mass spectrometric data, contributed to manuscript. Abhishek K. Jha oversaw development of PollyPhi Absolute, contributed to manuscript. Graeme F. Mason advised on design of PollyPhi Absolute, planned experiment design tests, wrote manuscript.

